# Simultaneous monitoring of the same animals with PIT-tags and sensor nodes causes no system interference

**DOI:** 10.1101/2020.03.05.978437

**Authors:** Simon P Ripperger, Niklas Duda, Alexander Kölpin, Gerald G Carter

## Abstract

Recent technological advances have multiplied the variety of biologgers used in wildlife research, particularly with small-bodied animals. Passive integrated transponders (PIT) have been used for decades to log visits of tagged animals at reader-equipped artificial feeders or roost boxes. More recently, novel miniaturized sensor nodes can collect data on social encounters among tagged individuals in any location. Combining these systems allows researchers to gather high-resolution tracking data on certain individuals from their long-term PIT-tagged animal populations. However, there can be a risk of interference among tracking systems. Here we tested whether placing an additional biologging sensor on top of a PIT-tag might attenuate the magnetic field reaching the PIT-tag and in turn hamper reading success of the radio-frequency identification (RFID) reader. We also evaluated data transmission by a digital sensor node in the presence of a magnetic field created by the RFID-antenna. The combination of this RFID-system and wireless biologging sensors works without error, suggesting that the simultaneous use of PIT-tags and other digital biologgers, e.g. miniaturized GPS-loggers, should also work together properly when communication channels do not overlap. The combination of long-term monitoring with PIT tags and short-term tracking with biologging sensor nodes creates exciting new opportunities to gather rich social data from individuals not present at RFID reader stations.

## Introduction

Biologging devices enable researchers to remotely gather information on the behaviour or physiology of free-ranging animals by means of animal-born tags. Passive integrated transponders (PIT-tags) provide a low-cost solution to identifying individuals either in the hand, or when they come near an antenna reader station. Using the latter method, called passive radio-frequency identification (passive RFID), researchers can create a log of visiting PIT-tagged animals at sleeping or feeding sites by mounting antenna readers around natural roosts such as tree holes (Patriquin et al. 2010; Toth et al. 2015) or by setting up reader-equipped artificial feeders or roost boxes (Aplin et al. 2015; Kerth and Reckardt 2003; Lopes et al. 2016; Nachev et al. 2017). The coil antenna creates a magnetic field, which triggers a PIT-tag when passing though the circular antenna to send back its identifying number. This method results in extensive datasets on individual use of any resources or locations that are equipped with antenna readers. While passive-RFID is an excellent low-cost method to gather extensive, highly standardized datasets in the long term, a general shortcoming is that data collection is restricted to the area near the antenna reader station.

A more complete picture of individual behaviour can be gained from increasingly powerful animal-borne biologging tags, such as GPS-loggers or proximity sensors. Recent technological advances have led to the miniaturization of these devices, creating new opportunities for studies of small animals (Ripperger et al. 2020). Another key advance in ‘next-generation’ biologgers is a diverse array of sensors such as accelerometers, magnetometers, or air pressure sensors, which autonomously collect and process data and give additional insights into the animal’s behaviour, performance, body posture, or flight height (O’Mara et al. 2019; Williams et al. 2017), which can be useful for studying foraging (Conenna et al. 2019; Egert-Berg et al. 2018; O’Mara et al. 2019) or social behaviour (Ripperger et al. 2019a; Ripperger et al. 2019b). However, due to the battery weight limitation and the rather high power demand of some tags, observation periods are often limited to only a few days or weeks. Still, these short-term studies can answer new questions beyond the reach of traditional methods. In particular, the combination of digital biologgers and long-term monitoring of generations of PIT-tagged individuals provides extraordinary opportunities to answer long-standing questions (Ripperger et al. 2020).

An important concern when combining tracking systems is that the simultaneous use of different technologies might cause interference or data loss in one or the other system. For example, RFID-reading success may depend upon an optimal orientation of the tag against the antenna plane, radio interference or the presence of metals within detection range (Aymes and Rives 2009). In bats, PIT-tags are often injected underneath the dorsal skin (Ellison et al. 2007; Kerth and Reckardt 2003; Neubaum et al. 2005; Rigby et al. 2011). So one important question is whether the magnetic field that reaches the PIT-tag is attenuated if another biologger is glued to the dorsal skin directly on top of the PIT-tag. A potential attenuation could hamper the identification of tagged bats since the PIT-tag draws the power for data transmission from the magnetic field of the coil antenna. Another possible but less likely concern is whether the magnetic field from the RFID-antenna will affect the functionality of the second digital device, since strong magnetic fields may impact MEMS (MicroElectroMechanical System)-based sensors. In this study, we test for interference between a popular RFID-system and sensor nodes from a novel wireless biologging network (Ripperger et al. 2020). We found that when a biologging sensor node was attached to the skin directly on top of a subcutaneous PIT tag, data transmission was not reduced in either system.

## Methods

We tested a widely used system for radio-frequency identification (RFID) consisting of subcutaneously implanted PIT tags (length 11mm; Trovan; Figure 1D) and a coil antenna (ca. 5.5 cm in diameter, Figure 1C) connected to a reader (Euro ID LID 665 Multi reader powered by a 12V motorcycle battery). For the second biologging device, we tested two versions of recently developed sensor nodes (part of the BATS wireless biologging network (Duda et al. 2018; Ripperger et al. 2020); see Figure 1). These sensor nodes contain an accelerometer, a magnetometer and an air pressure sensor which continuously sampled data and wirelessly transmitted them to a receiver unit. The first tested version of the sensor nodes was built from populated flex substrate (22mm × 14mm × 1mm; length × width × height, without antenna; Figure 1A, B) connected to a LiPo battery as previously deployed in bat research (Ripperger et al. 2019a; Ripperger et al. 2019b). To mimic larger biologgers such as GPS-tags, the second version was built from thicker FR4, a common material for printed circuit boards (33mm × 13mm × 3mm; without antenna or plugs; Figure 1A).

**Figure 1:**
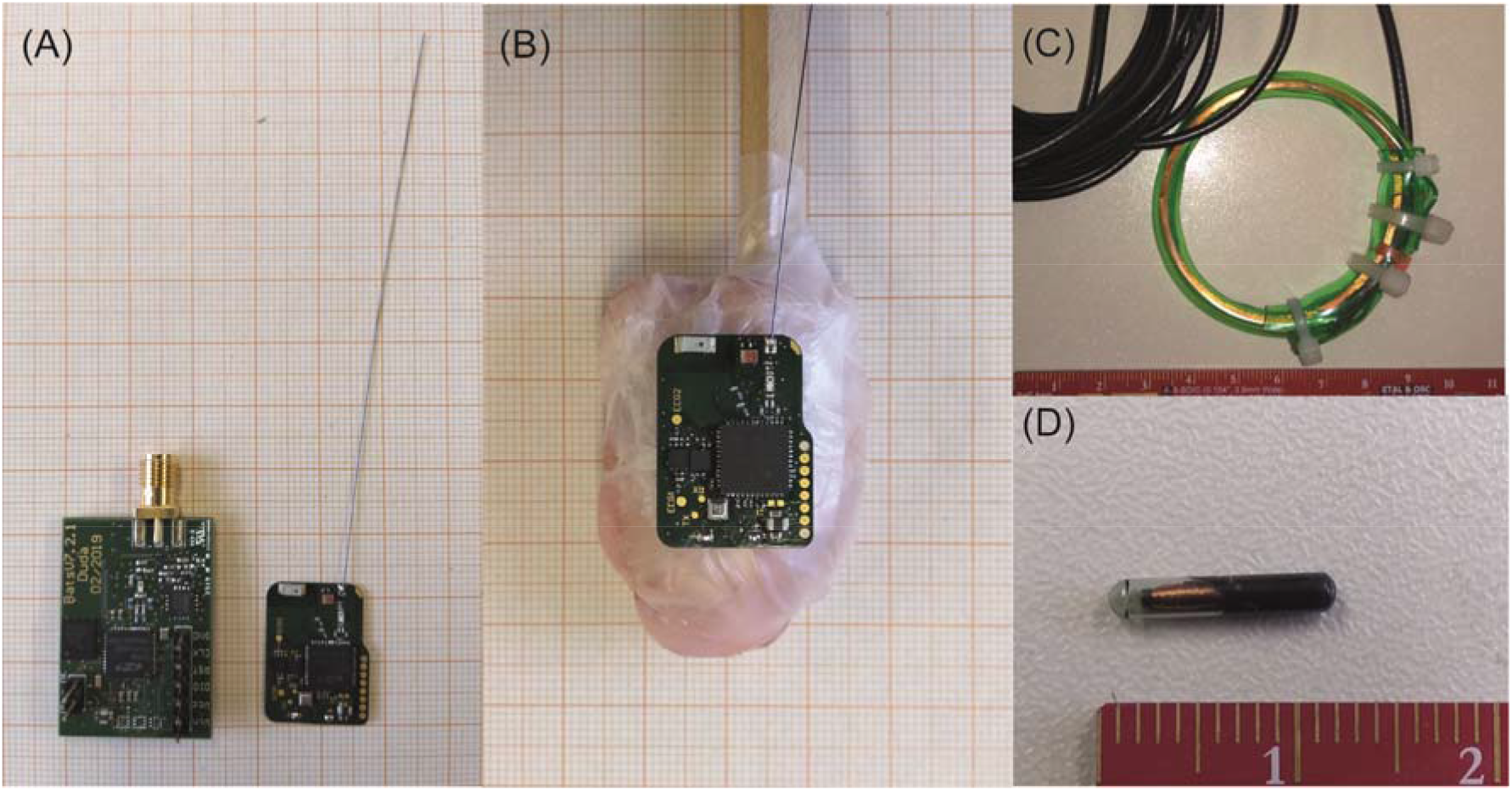
Test hardware and bat dummy. (A) larger FR4 and smaller flex sensor nodes, (B) test dummy made from chicken meat with subcutaneously injected PIT tag and flex sensor node attached (inactive, without battery), (C) coil antenna of the RFID system, (D) Trovan PIT tag. (A, B) smallest squares on grid paper are 1 mm^2^; (C, D) ruler shows centimeters.

To simulate a medium-sized bat, we first created a test dummy from 15 g meat off the bone of a raw chicken wing (organically raised chicken meat) wrapped in parafilm in an ellipsoid shape (Figure 1B; the sensor node is placed on top for a comparison in size; during the experiments, the sensor node was connected to a LiPo-battery and wrapped in parafilm). We ‘subcutaneously’ injected a PIT tag in the dorsal centre of the dummy, mounted on a ca. 15 cm wooden stick. For the following tests, we held the dummy by the stick and moved it back and forth through the antenna of the reader at a speed similar to a bat crawling through the entrance/exit of a bat box. The RFID data were recorded on a PC connected to the antenna reader using the software Dorset ID V804. Data from the sensor nodes were received via a custom-made radio-frontend connected with USB to a PC by a script written in Python 3.7.

To identify whether the sensor node decreased the RFID success rate of the underlying PIT tag, we tested the dummy with the subcutaneously injected PIT tag under three conditions: (a) no sensor (control), (b) flex sensor node, (c) the thicker FR4 sensor node, mounted on top. In each condition, we moved the dummy back and forth 50 times through the coil antenna (i.e. 100 passages). We counted the number of passages registered by the RFID software.

We also tested if the magnetic field of the coil antenna led to reduced communication between the sensor node and its receiver in the form of ‘package loss’. Briefly, data from all sensors (e.g. acceleration, magnet field and air pressure) is transmitted in a uniquely labelled package of 27 bytes. We measured package loss while moving the flex sensor-equipped dummy back and forth through the active coil antenna as described above and while it was held stationary in the centre of the coil antenna. As a control, we repeated this procedure with an inactive coil antenna (disconnected from the power source). Finally, to evaluate how much the magnetic field of the RFID antenna affects the magnetometer on the sensor, we also plotted the magnetometer data.

## Results

Radio-frequency identification of the PIT tag injected in the dummy was successful in 100 of 100 cases in all three conditions (i.e. no sensor node, the thin flex sensor node, and the thicker FR4 sensor node, placed on top of the PIT tag). The magnetic field of the coil antenna of the PIT tag reader did not cause packet loss (packet loss with an inactive coil antenna: 0.067%, n = 10436 packets; packet loss when moving through active coil antenna: 0.039%, n = 17,748 packets; packet loss when stationary in active coil antenna: 0.059%, n = 6,777 packets). The magnetic field generated by the coil antenna of the PIT tag reader added only ca. 0.07 Gauss to the overall signal level of ca. −0.41 Gauss during every passage through the center of the reader coil, which was marked by a short peak in magnetic flux (Figure 2).

**Figure 2:**
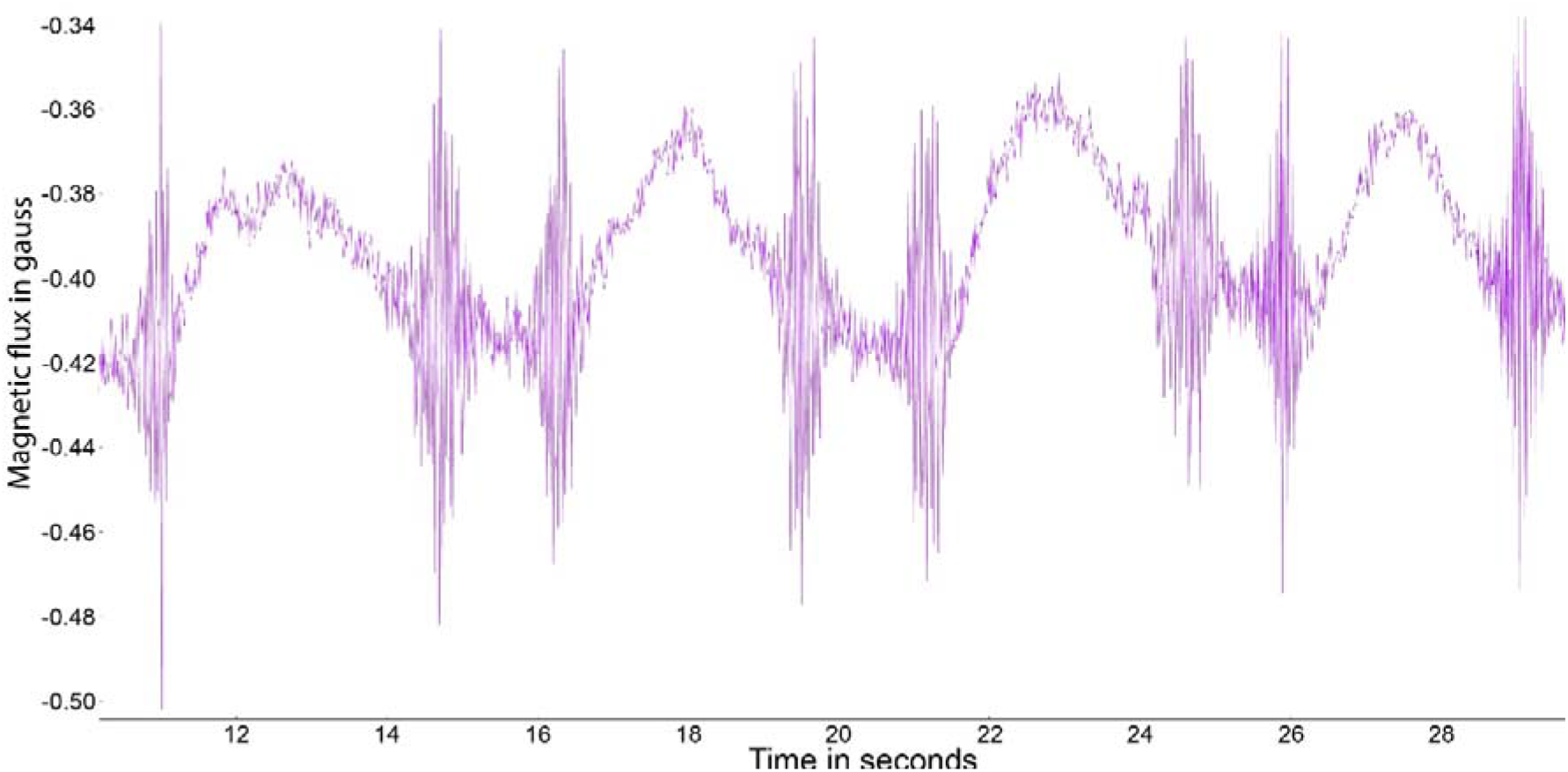
Magnetometer-generated data while moving the sensor tag back and forth through the RFID reader coil. Pronounced peaks mark the passage through the center of the coil.

## Discussion

The combination of the Trovan RFID-system and the BATS wireless biologging network works without error, and systems using similar mechanisms should also work properly together. Data transmission was normal in the RFID system and using the sensor nodes made from either the thin or thicker substrate. The only impact we discovered was that the magnetometer on the sensor node was affected as expected while in range of the RFID reader coil’s magnetic field (Figure 2). Magnetic field strength measurements, to evaluate animal body posture for instance (Williams et al. 2017), should therefore be treated with caution when evaluating data from animal-borne tags near PIT-tag readers. However, a positive byproduct of the occurrence of this distinct signal in the magnetometer-generated data is that these events could be used to synchronise timers between the two systems. High-power RFID coils also might have an impact on other MEMS-based sensors like the accelerometer or air pressure sensor, but the 0.07 Gauss measured in our setup were too weak to cause such problems.

In general, it is vital to operate animal tracking and biologging systems on different radio frequencies to avoid interference. The BATS system operates on frequency bands at 868 MHz (Europe/Asia) or 915 MHz (Americas) and 2.4 GHz (worldwide). Most PIT-tag systems for radio-frequency identification of animals operate at 125-150 kHz or 13.56 MHz (Bonter and Bridge 2011). These frequencies are so far apart that interference can be ruled out. However, there are RFID systems available that operate at 868 or 915MHz. When using such systems in combination with other systems also operating in the same band, it is necessary to either specify different frequency channels that do not overlap or - if the system allows for such customization - to use channel access control mechanism to prevent interference. The frequencies of background signals from global navigation satellite systems (GNSS) like GPS, Galileo or Glonass, should not interfere with other commercial wildlife tracking devices, since those are exclusively reserved for GNSS by global regulations (frequencies between 1160 MHz and 1590 MHz (Gao and Enge 2012)).

With transmission frequencies of different biologger systems far enough apart, there is no risk of interference of the transmitted signals. The combination of long-term monitoring with PIT tags and short-term tracking with biologging sensor nodes creates exciting new opportunities to gather rich data from individuals when they are not present at RFID reader stations, e.g. Ripperger et al. (2020).

## Acknowledgements

We thank G Kerth and J van Schaik for providing the PIT-tags and the RFID reader system. This study was funded by Deutsche Forschungsgemeinschaft (DFG FOR-1508).

## Conflict of interest

The authors declare that no conflict of interest exists.

## Notes

### Competing Interest Statement

The authors have declared no competing interest.

